# Parsing the Functions of Immediate Early Proteins in HCMV Infection Outcome

**DOI:** 10.1101/2025.09.19.676120

**Authors:** Yaarit Kitsberg, Aharon Nachshon, Noam Stern-Ginossar, Michal Schwartz

## Abstract

Human cytomegalovirus (HCMV) is a prevalent pathogen of the herpesvirus family, infecting most of the human population worldwide. Like all herpesviruses, HCMV can establish a latent infection that persists throughout the lifetime of the host. The HCMV immediate early (IE) proteins, IE1 and IE2, are viewed as master regulators of HCMV infection and are commonly assumed to play pivotal roles in regulating the balance between latent and lytic infection, as their repression is a hallmark of latency. However, it is still unclear whether their expression can indeed determine the establishment of productive infection and what functions, either related to viral gene expression or to cellular pathways, are involved in this activity. Using THP1 monocytes, ectopically expressing the HCMV receptor, PDGFRα to boost viral entry, we show that overexpression of either IE1 or IE2 significantly enhances productive infection, illustrating their critical role in determining infection outcome. Mechanistically, we show IE2 drives expression of the viral early genes at early stages of infection, whereas IE1 acts more broadly to enhance global viral gene expression. We further show that from the many functions of IE1, its ability to promote lytic infection is mainly linked to the disruption of PML nuclear bodies. Importantly, induction of either IE1 or IE2 expression in latently infected cells enhances viral reactivation, with IE1-mediated PML representing a central mechanism. Taken together, our findings elucidate the distinct and complementary roles of IE1 and IE2 in overcoming barriers to productive infection and reactivation.

## Introduction

Human cytomegalovirus (HCMV) is a prevalent pathogen of the herpesvirus family, infecting the majority of the human population worldwide. Like all herpesviruses, HCMV can establish a latent infection that persists throughout the lifetime of the host. Reactivation from latency in immunocompromised individuals, such as transplant recipients and HIV patients, can result in severe morbidity and mortality and continues to be a serious healthcare issue ^1,2^.

Latent infection is defined by maintenance of the viral genome in the absence of viral progeny production, with the ability to reactivate full lytic infection. HCMV latency has been studied and characterized in progenitor cells of the myeloid lineage, such as haematopoietic stem cells (HSCs), granulocyte–macrophage progenitor and blood monocytes ^3–5^. On the other hand, terminally differentiated myeloid cells, e.g. macrophages and dendritic cells are more prone to lytic infection and furthermore, differentiation of infected progenitor cells, such as monocytes, may cause reactivation of the virus.

Lytic infection of HCMV starts with transcription of viral immediate early (IE) genes from the major immediate-early promoter (MIEP). The two major proteins encoded from the MIE transcript, namely IE1 (also IE72 encoded by UL123) and IE2 (also IE86 encoded by UL122), are products of alternative splicing of a single locus composed of 5 exons. The canonical IE1 is expressed from exons 1–4, while IE2 is expressed from exons 1–3 and 5^6,7^. IE2 is a strong transactivator of viral early gene expression ^8–11^ and is indispensable for viral early gene expression and viral growth ^11,12^. IE1 also has an important role in activation of viral gene expression, and its knockout attenuates (but does not completely eliminate) viral replication ^13–16^. IE1 is a multi-functional protein ^17^; It modulates chromatinization of the viral genome via association with HDAC3. Indeed, treatment with histone deacetylase (HDAC) inhibitors rescued the growth defect of the IE1-deletion virus ^18^. In addition, IE1 disperses the antiviral nuclear bodies termed promyelocytic leukaemia (PML) bodies ^19,20^ that play a part in the defense against viruses ^21^, and specifically were shown to restrict HCMV infection ^22^. Finally, IE1 was shown to dampen the response to interferon by blocking downstream signaling through interaction with STAT proteins ^23,24^. All these functions contribute to its role in supporting activation of viral gene expression.

In contrast to lytic infection, latent infection is characterized by chromatin-mediated repression of the MIEP ^25–28^. While viral gene expression is globally repressed during latency^29–31^, IE genes are subject to additional distinct levels of repression ^29,32^. Nevertheless, the contribution of the specific IE gene repression to the onset of latent infection is far from understood. If repression of IE gene expression is a major factor in establishment of latency, one would expect that increasing their expression would enable the establishment of productive infection. Indeed, productive infection of monocyte-derived macrophages (MDMs), as opposed to latent infection, is characterized by robust expression of IE1 and IE2 and overexpression of either IE1 or IE2 results in significant increase in the percentage of cells that succumb to productive infection ^33^, meaning these genes have a major role in determining HCMV infection outcome. However, in monocytes, which largely support latent infection, overexpression of IE genes was not sufficient to induce productive infection ^34^, pointing to potentially additional factors regulating latency.

Here, we recapitulated these observations that overexpression of IE genes in THP1 monocytes have very weak effects on productive infection. Nevertheless, when increasing viral entry by overexpression of the HCMV receptor, PDGFRα, we found that overexpression of either IE1 or IE2 robustly promotes productive infection in monocytes. We found that IE1 globally enhances viral gene expression and drives productive infection mainly through disruption of PML nuclear bodies. IE2 on the other hand, increases infection likely by specifically upregulating the expression of early viral genes. Strikingly, we further reveal that when inducing the expression of either IE1 or IE2 in latent cells, viral reactivation takes place, in support of their critical role in dictating infection outcome. Overall, our data demonstrate the important role of IE1 and IE2 in regulating latent infection and reactivation, through modulation of PML nuclear bodies and regulation of viral gene expression, respectively.

## Results

### Ectopic expression of IE genes promotes productive infection in monocytes upon enhanced viral entry

To examine whether limited expression of the immediate-early proteins IE1 and IE2 is a barrier for productive infection in monocytes, which largely support latent infection, we overexpressed IE1 and IE2 in THP1 monocytic cells (Fig. S1a). To minimize potential artifacts from ectopic expression of these genes, we utilized a Doxycycline (Dox)-inducible system and induced expression only at the time of infection. Infection was performed with the TB40/E strain expressing GFP ^35^, which allows quantification of productive infection. Analysis at three days post-infection (dpi), revealed that overexpression of either IE1 or IE2 had only a minor effect on productive infection of THP1 cells (Fig. 1a), in line with previous findings ^34^. This points to a potentially fundamental difference in infection between monocytes and macrophages, since overexpression of either IE1 or IE2 does promote productive infection in macrophages, illustrating their levels of expression serve as a barrier for productive infection^33^. We thus tested whether IE proteins can promote lytic infection also in primary fibroblasts, which are common models for lytic HCMV infection, and found that when infected at low MOI, overexpression of either one of the IE proteins enhances productive infection in fibroblasts (Fig. 1b). Thus, illustrating that IE1 and IE2 levels of expression indeed serve as an important barrier for HCMV productive infection.

**Fig. 1.**
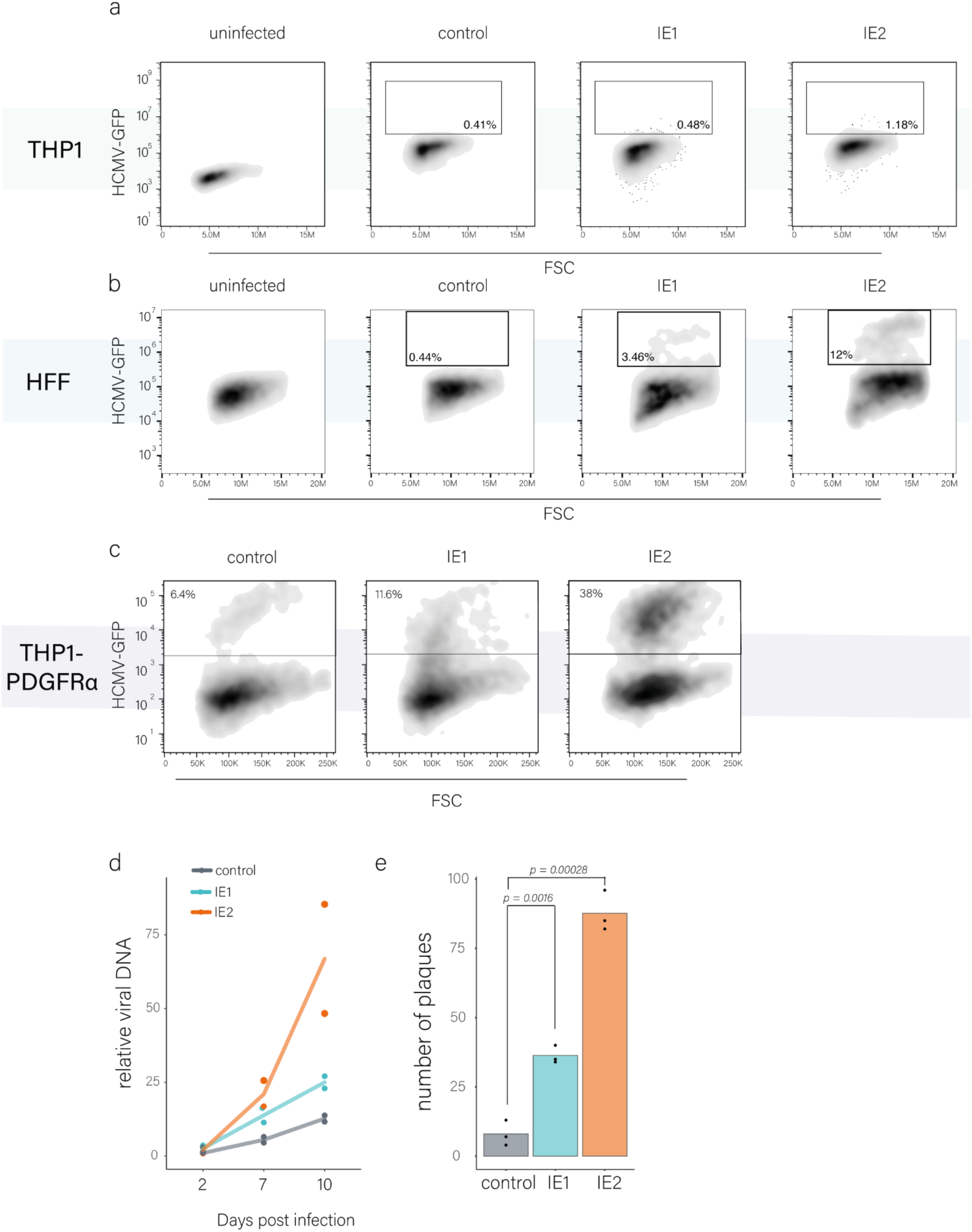
(a). Flow cytometry analysis of THP1 cells induced to overexpress control, IE1, or IE2 genes using a Tet-On inducible system at the time of infection. Cells were analyzed 3 days post-infection (d.p.i.). (b). Flow cytometry analysis of HFF cells induced to overexpress control, IE1, or IE2 genes using a Tet-On inducible system at the time of infection. Cells were analyzed 3 days post-infection. (c). Flow cytometry analysis of THP1-PDGFRα cells overexpressing control, IE1 or IE2 proteins at the time of infection with HCMV-GFP using the Tet-On inducible systems. Cells were analyzed at 3 d.p.i. (d). Viral DNA replication levels of THP1-PDGFRa overexpressing control, IE1 or IE2 at 2, 7 and 10 days post HCMV infection. Analysis was performed using qPCR. (e). Measurements of HCMV progeny in supernatants collected from infected THP1-PDGFRα overexpressing control, IE1 or IE2 at the time of infection. The supernatant was collected at 10 d.p.i. p-value was calculated using a two-sided student t-test. n=3.

We recently showed that entry of HCMV into monocytes is substantially less efficient than in macrophages, and that ectopic expression of one of the HCMV entry receptors, PDGFRα or THBD, in THP1 cells leads to partially productive infection of monocytes ^36^. To test if limited virus entry is a significant barrier for infection in monocytes that masks IE1 and IE2 potential importance, we next tested the effect of IE1 or IE2 overexpression in THP1 ectopically expressing PDGFRα (THP1-PDGFRα). Overexpression of either IE1 or IE2 significantly and robustly increased the percentage of productively infected (GFP-bright) cells compared to the control cells, in which mCherry was overexpressed (Fig. 1c). This increase was associated with enhanced viral DNA replication (Fig. 1d) as well as increased viral progeny (Fig. 1e), although the production of infectious progeny is low in these cells, due to cis inhibitory effects mediated by the constitutive expression of PDGFRα ^36^.

Overall, these results demonstrate that the expression levels of IE proteins are key determinants of productive infection in diverse cell types including monocytes, indicating they constitute a significant barrier for HCMV lytic infection.

**Fig. S1.**
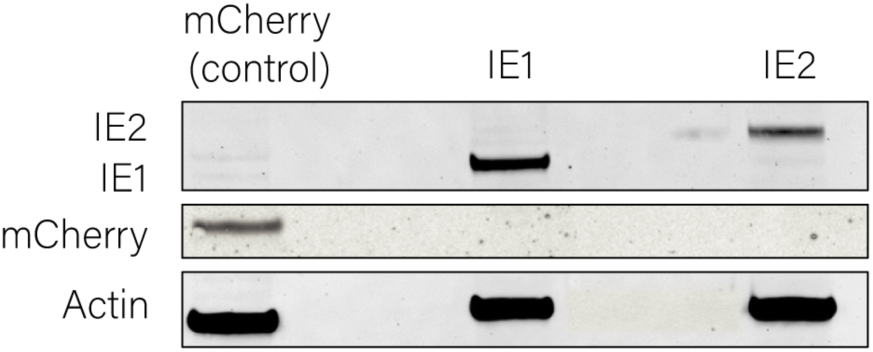
Western blot analysis of THP1 overexpressing mCherry control, IE1 and IE2 24 hours after Dox induction.

### IE1 globally enhances viral gene expression, while IE2 specifically enhances expression of early viral genes

Both IE proteins are multifunctional, promoting viral gene expression while modulating diverse cellular pathways ^17,32^, however, it remains unclear which of these activities are critical for determining infection outcome. To examine this, we first characterized the influence of IE1 and IE2 on viral gene expression as well as on cellular pathways in monocytes, by performing RNA-seq on THP1-PDGFRα cells ectopically expressing either protein, in uninfected cells as well as at early stage of infection (12 hours post-infection). We initially verified that, in the absence of Doxycycline induction, expression of IE1 and IE2 was negligible (Fig. S2a), and no transcriptional changes compared to the control cells were observed (Fig. S2b), excluding the possibility of leakage in IE gene expression. Quantification of viral transcript load, as measured by the percentage of viral reads out of total reads, demonstrated that overexpression of IE1 resulted in increased viral gene expression while overexpression of IE2 did not dramatically change the global levels of viral genes (Fig. 2a). To investigate IE1 and IE2 effects on specific viral genes, we quantified the relative expression of distinct temporal classes of viral genes ^29^. We found that overexpression of IE1, while broadly enhancing viral gene expression (Fig. 2a), does not affect specific viral gene classes (Fig. 2b), or viral genes (Fig. 2c). This indicates that IE1 globally activates viral gene expression, likely by reducing repression imposed on the viral genome ^18^.

**Fig. 2.**
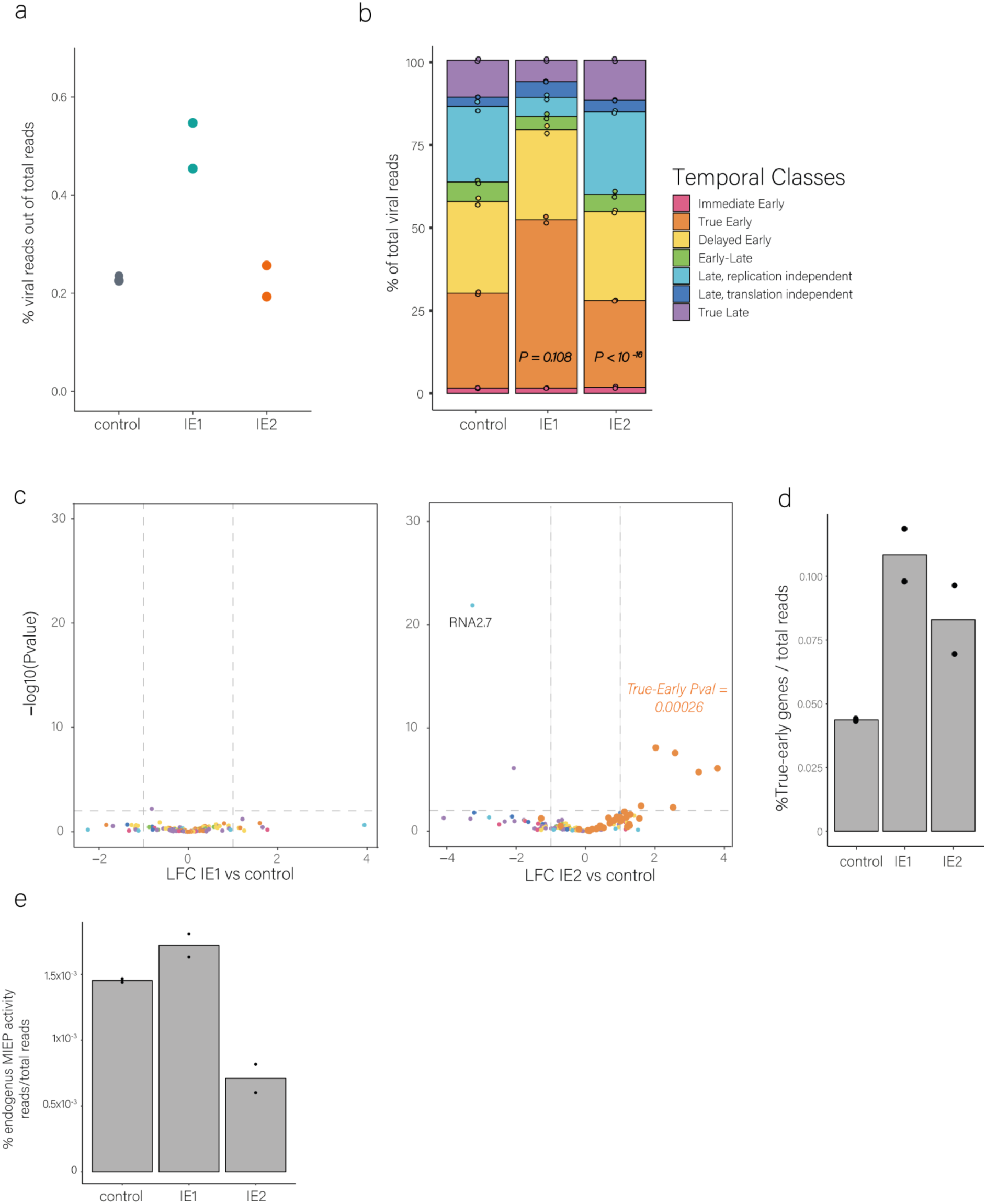
(a). Percentage of viral gene expression out of total gene expression in infected THP1-PDGFRa cells overexpressing IE1, IE2 and control. Cells were collected for RNA-seq at 12 hours post-infection. (b). Percentage of reads corresponding to specific viral temporal classes out of total viral gene expression in THP1-PDGFRa cells overexpressing IE1, IE2 and control. Statistics were performed using the Hypergeometric enrichment test. *UL122* and *UL123* transcripts were omitted from the calculation (c). Differential expression analysis of viral genes in THP1-PDGFRa overexpressing IE1 (left) or IE2 (right) versus the control. Cells were collected at 12 hours post-infection. Viral genes were marked according to their temporal classes. Enrichment statistics performed using Hypergeometric enrichment test. *UL122* and *UL123* transcripts were omitted from the calculation (d). Percentage of the early set of genes (True-early) out of total gene expression in THP1-PDGFRa cells overexpressing IE1, IE2 and control. (e). MIEP activity of cells overexpressing IE1, IE2 or control normalized by total host reads. The activity was measured by summing reads on the 5’ and 3’ UTRs of the IE transcript.

IE2 on the other end, significantly increased the relative expression of true-early viral genes compared to other viral transcripts (Fig. 2b,c and Table S1) and also compared to total transcripts (Fig. 2d), indicating an absolute increase in the transcription of true-early genes^10,11,15^.

Surprisingly, IE2 also seems to downregulate the expression of late replication-independent genes (TC6), but examinations of individual genes showed IE2 expression specifically reduced RNA2.7 expression, (Fig. 2 b,c). This effect seems specific as it was also observed when IE2 was expressed prior to infection (Fig. S2c). This suggests an additional role for IE2, which on top of initiating the expression of true early genes, it also specifically suppresses the expression of RNA2.7, which is the most abundant viral transcript in HCMV infected cells ^37,38^.

Our RNA-seq data also enabled us to examine the autoregulatory effects of IE proteins on the MIEP by measuring expression from the 3ʹ and 5ʹ UTRs of *UL123* and *UL122*, which are not part of the overexpressed transgenes. Overexpression of IE1 modestly increased transcription from the MIEP (as measured by the abundance of reads mapped to the 5’ and 3’ UTR), which is possibly as part of the global enhancement of viral gene expression, whereas in agreement with earlier works ^8,39^, IE2 reduced transcription from the MIEP (Fig. 2e).

**Fig. S2.**
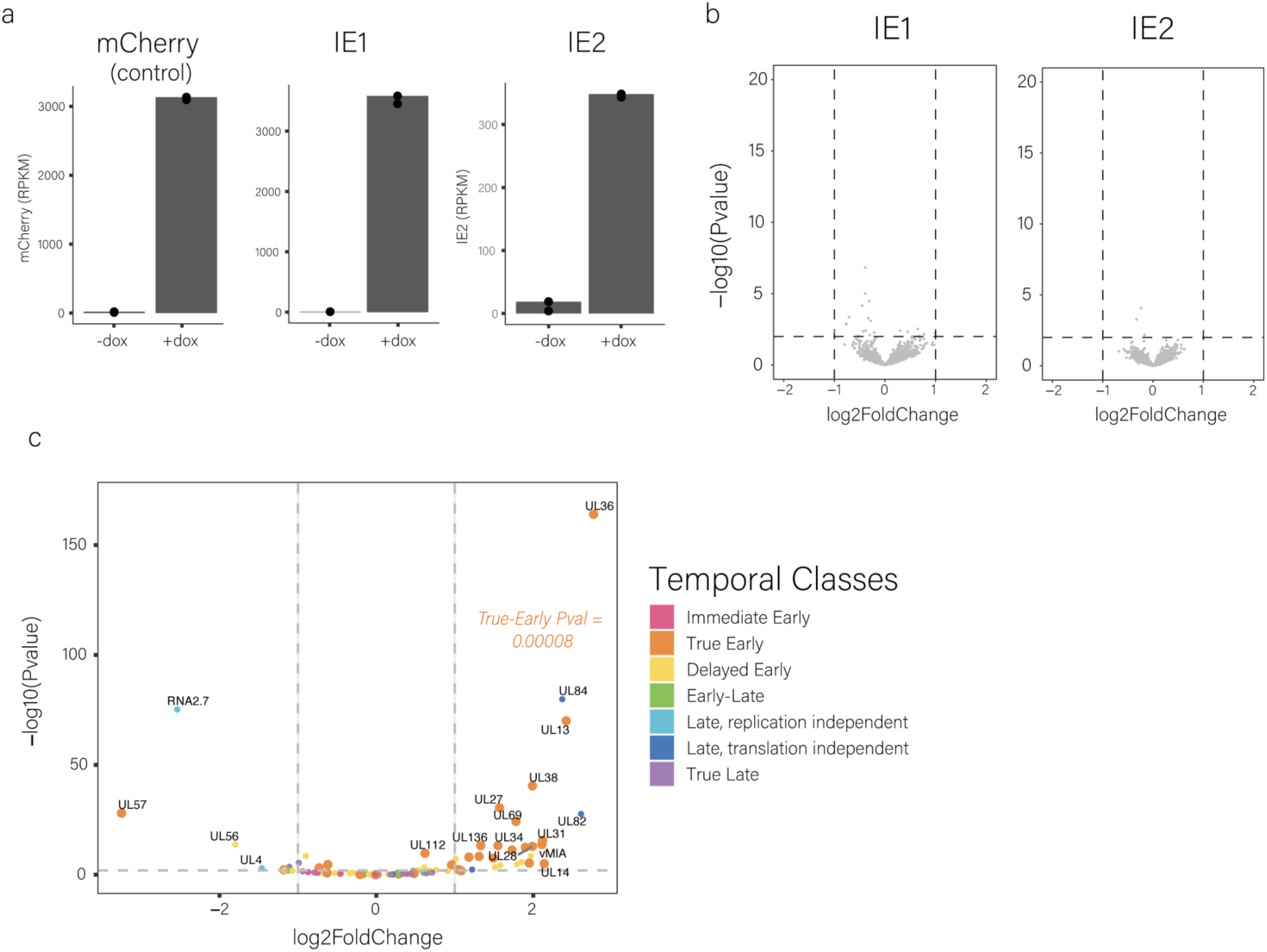
(a). Expression levels of mCherry (control), IE1 and IE2 in THP1-PDGFRα cells, either untreated or treated with Dox for 24 hours, as measured by RNA-seq. (b). Volcano plot showing differentially expressed genes in THP1-PDGFRα cells without Dox induction of IE1 and IE2 compared to the control. (c). Differential expression analysis of viral genes in THP1-PDGFRa overexpressing IE2 versus control. Cells were induced to express IE2 or control for 24 h followed by HCMV infection. Samples were collected at 12 hours post-infection. Viral genes were marked according to their temporal classes. Enrichment statistics was performed using the Hypergeometric enrichment test. *UL122* and *UL123* transcripts were omitted from the calculation.

### Distinct Roles of IE1 and IE2 in Innate Immunity and Cell Cycle Regulation

We also analyzed the transcriptome data to gain an unbiased view on the cellular pathways that are affected by IE1 and IE2 in monocytes. The most striking effect of IE1 overexpression was reduction in pathways of interferon response, likely due to its binding to STAT2 and inhibition of ISG induction by ISGF3 ^24,40^ (Fig. 3 a,b, c and Table S2). Additional pathways such as the NFkB and other inflammatory pathways were also reduced but much less in uninfected cells, pointing to involvement of additional viral factors in this process. On the other hand, other pathways related to cell cycle progression and mTOR signaling were significantly reduced by IE1 overexpression, but much less so in the context of infection (Fig 3a).

**Fig. 3.**
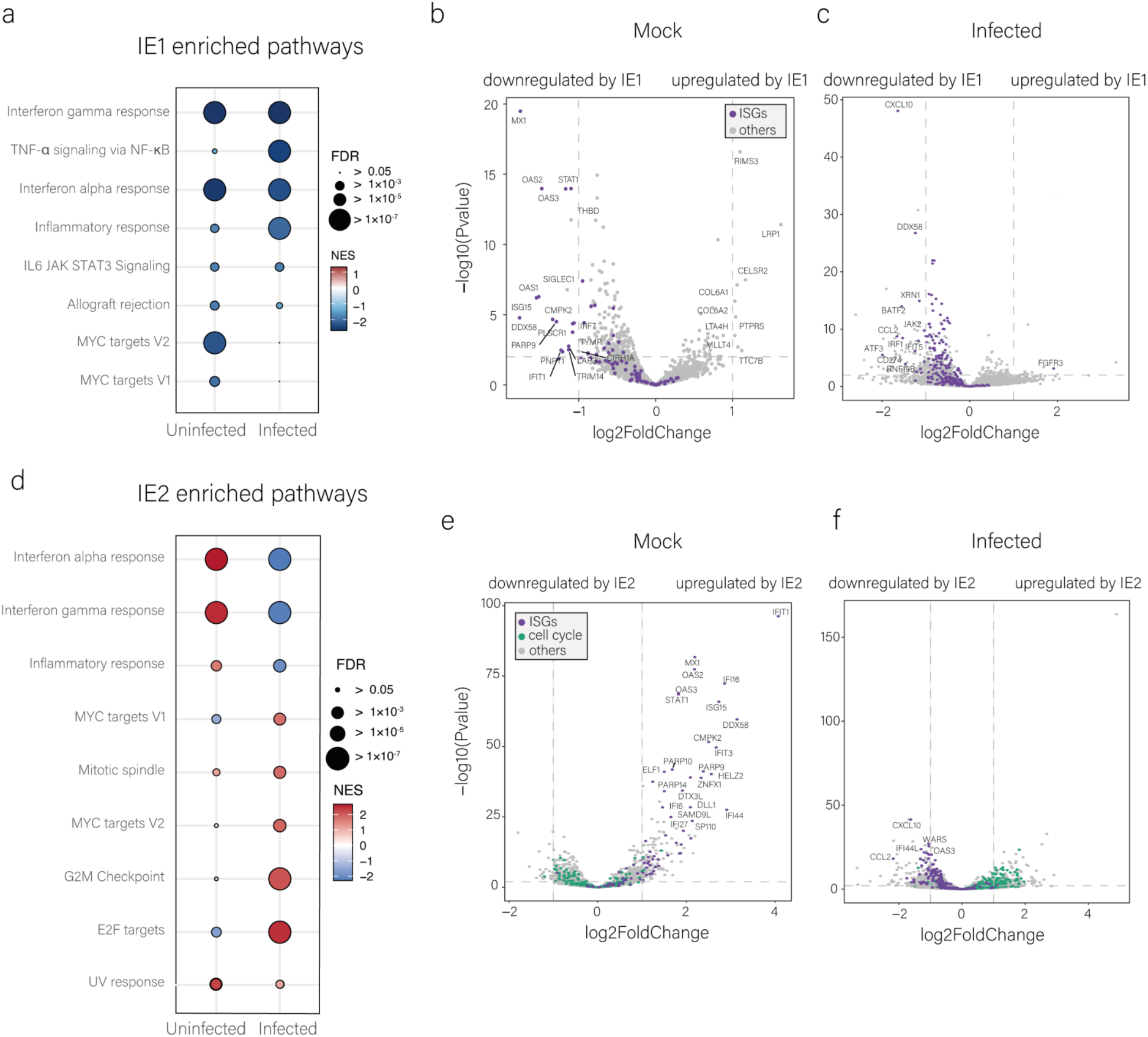
RNA-seq analysis of uninfected and infected cells (12 hpi) overexpressing IE1 or IE2 compared to control. a, d Summary of Hallmark pathway enrichment analysis of differentially expressed genes upon ectopic expression of IE1 (a) or IE2 (d) for 24 h in uninfected cells (mock) or in cells induced to express IE1 (a) or IE2 (d) at the time of infection and collected at 12 hours post-infection. FDR, false discovery rate; NES, normalized enrichment score. b and c, Volcano plots showing the differentially expressed genes in IE1 overexpression compared to the control in uninfected (24 h induction, b) and infected cells (12 hpi, c). Purple dots indicate interferon-stimulated genes. e and f, Volcano plots showing the differentially expressed genes in IE1 overexpression compared to the control in uninfected (24 h induction, e) and infected cells (12 hpi, f). Purple dots indicate interferon-stimulated genes. Green dots indicate cell-cycle related genes.

Similarly, the strongest enrichment detected when IE2 is expressed was seen for pathways of interferon response, however strikingly, the effects were opposite in uninfected and infected cells (Fig. 3 d, e, f and Table S2). Ectopic expression of IE2 in THP1 cells led to a strong induction of interferon response in uninfected cells (Fig. 3d, 3e, S3), and in contrast to downregulation of interferon response in infected cells (Fig. 3 d,f). We postulate that the induction of IFN-stimulated genes in uninfected cells overexpressing IE2 may be related to IE2-mediated induction of DNA damage ^41^. DNA damage was shown to directly induce the interferon response ^42^, potentially explaining IE2 driven IFN induction. IE2 is known to induce genes required for host cell DNA synthesis and to concurrently inhibit cellular DNA synthesis ^43^ and indeed there is strong upregulation of cell cycle related pathways when it is overexpressed (Fig. 3d). This effect is more pronounced in the context of infection, perhaps due to involvement of additional viral proteins or to the lack of deleterious effects of IE2 outside of the context of infection.

**Fig. S3.**
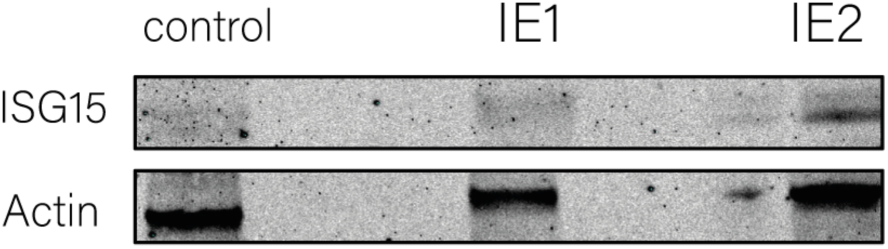
Western blot analysis of THP1 overexpressing mCherry control, IE1 and IE2 24 hours after Dox induction. The membrane was stained with the innate immunity related gene ISG15 and Actin as a housekeeping protein.

### Temporal Dynamics of IE Gene Effects on Viral Infection

Transcriptome analysis of IE gene overexpression in both infected and uninfected cells revealed distinct functions of IE1 and IE2. IE1 broadly enhances viral gene expression, consistent with a role in relieving repression on the viral genome, and concurrently dampens the interferon response. IE2 specifically and directly activates transcription of viral early genes ^10,11,15^, and also influences cell cycle and interferon response pathways. To better understand the potential roles of these different effects in affecting infection outcome, we investigated whether the ability of IE1 and IE2 to promote productive infection is restricted to a particular time window. To address this, we induced IE1 or IE2 expression at different timepoints: at the time of the infection (0 hours post-infection, hpi), 5hpi or 24hpi, and measured infection at 3 dpi. Additionally, due to the different effects of these proteins in uninfected cells, we also induced the IE proteins 24 hours before infection (−24hpi),

When overexpression is induced prior to infection (−24h), IE1 led to an even stronger effect on productive infection than when induced at the time of infection, while the effect of IE2 diminished (Fig. 4a and S4a-c). These results fit the opposing effects on innate immunity pathways in uninfected cells, IE1 dampens while IE2 enhances, effects which likely contribute to the effect on infection, in line with previous reports showing that induction or reduction of ISG expression prior to infection decreased or increased productive infection, respectively ^44,45^. IE1 significantly enhances productive infection also when induced at the time of infection, but not when it is induced at later time points. IE2 on the other hand, significantly enhances productive infection when induced at any of the time points tested during infection. These results demonstrate that IE1 is able to promote productive infection within a short time window at the onset of infection, whereas the effect of IE2 is more prolonged.

**Fig. 4.**
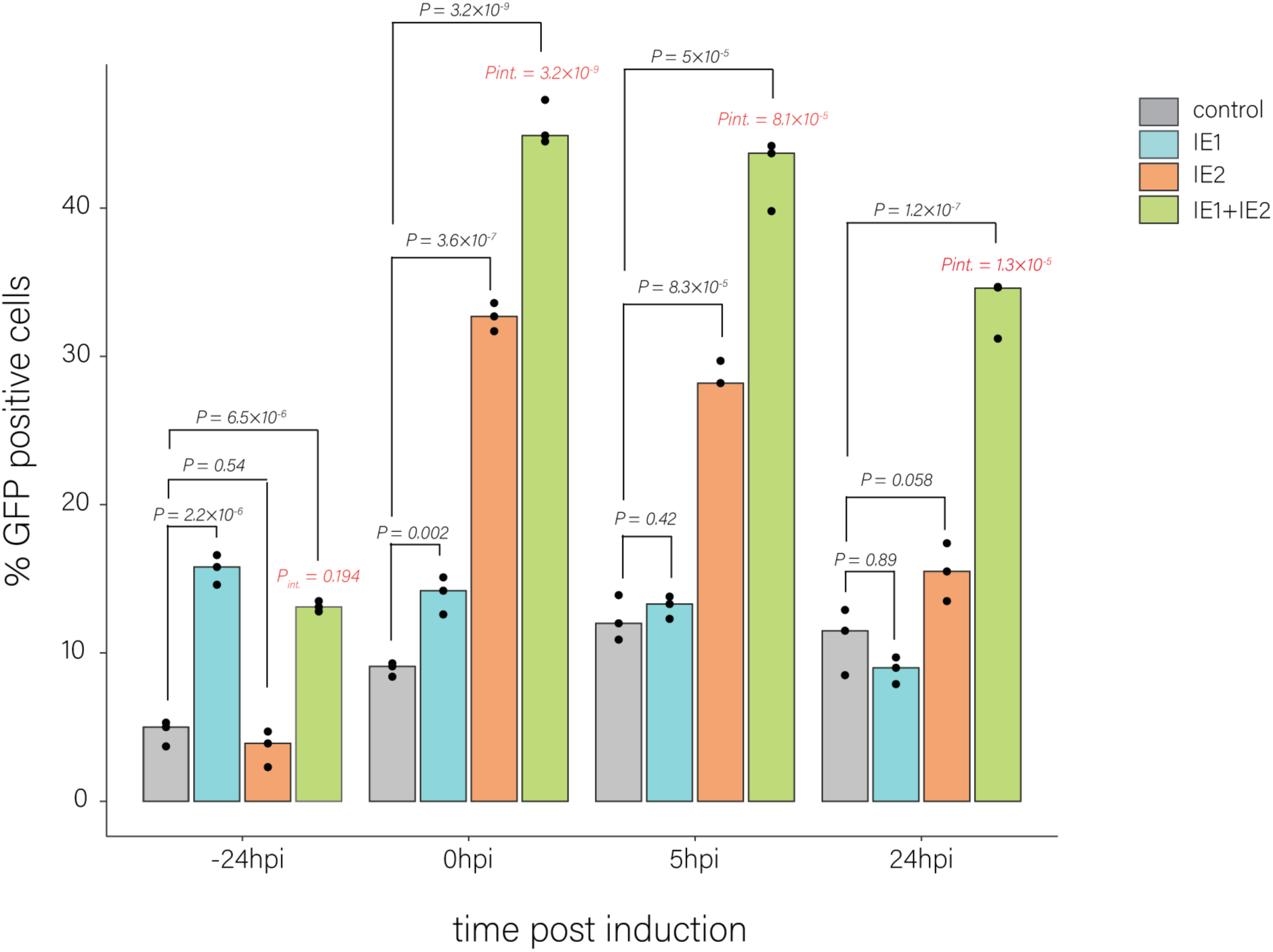
(a). Flow cytometry analysis of infected THP1-PDGFRa cells that were induced to express IE1, IE2 or control at 24 hours prior to infection (−24hpi), at the time of infection (0hpi), 5hpi and 24hpi. Cells were analyzed at 3 d.p.i. p-value was calculated using a two-sided student t-test and ANOVA for interaction. P_int_ indicates the P-value of the interaction. n=3.

We further examined the combined effect of IE1 and IE2 on infection at different times of induction. When these genes are induced prior to infection, the combined effect is similar to the effect of IE1 alone. When induced at the time of infection or subsequently, the combined effect of IE1 and IE2 on infection exceeded the sum of their individual effects, indicating a synergistic interaction (Fig. 4), suggesting a concerted function that enhances their individual activities. This synergism may be due to either the distinct complementary functions of the two proteins, or from one protein enhancing the function of the second protein ^9,10^.

**Fig. S4.**
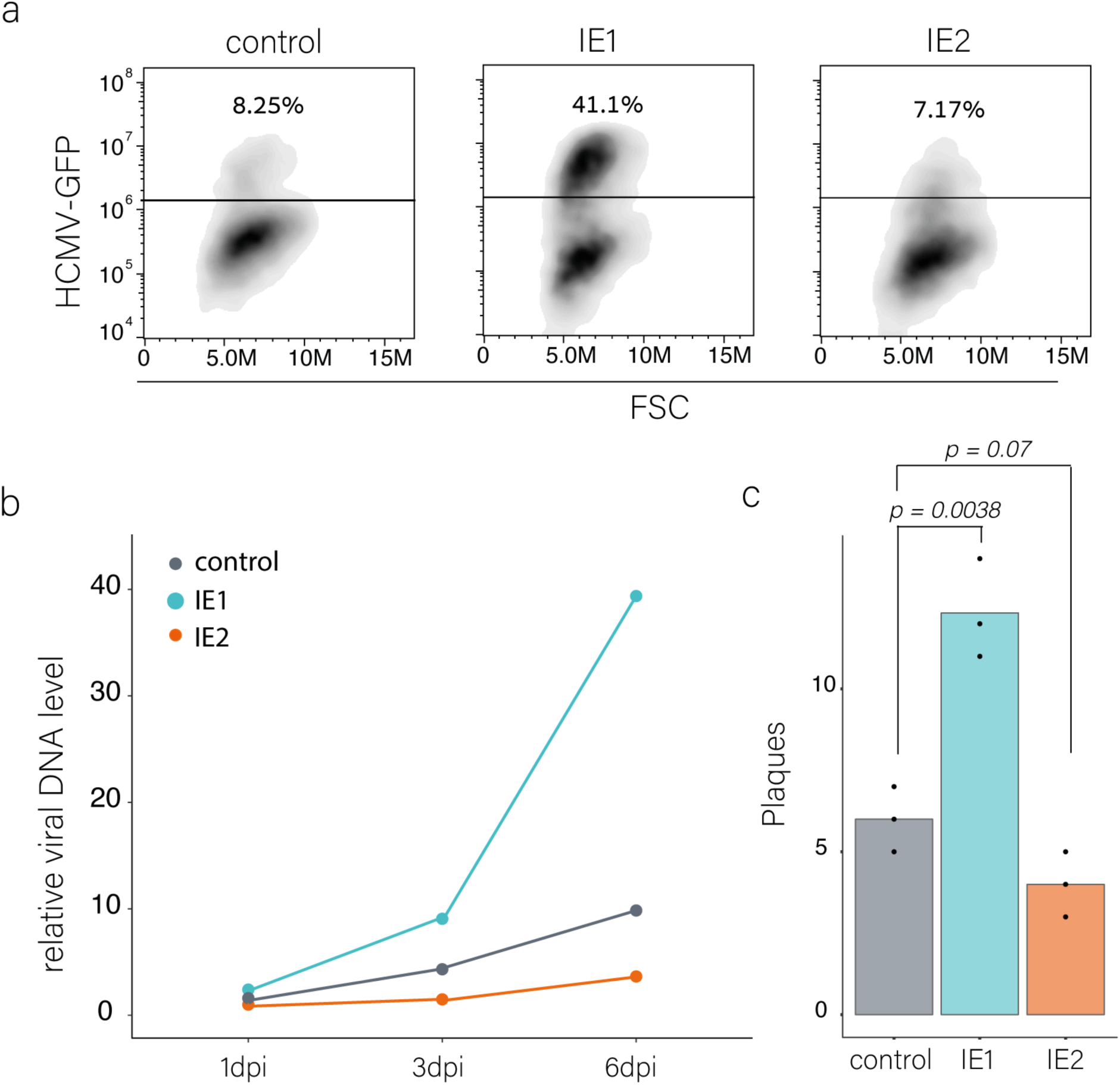
(a). FACS analysis of THP1-PDGFRa ectopically expressing control, IE1 or IE2 proteins 24 hours before infection using the Tet-On inducible systems. Cells were analyzed at 3 d.p.i. (b). Viral DNA replication of THP1-PDGFRa expressing control, IE1 or IE2 at 1, 3 and 6 d.p.i. Ectopic expression was induced using the Tet-On inducible systems 24 hours before infection. Analysis was performed using qPCR. (c). Measurements of infectious virus in supernatants collected from infected THP1-PDGFRa overexpressing control, IE1 or IE2 at 10 d.p.i. Induction using the Tet-On inducible systems was done 24 hours before infection. p-value was calculated using a two-sided student t-test. n=3.

### IE1 promotes productive infection predominantly by inhibiting PML association

IE1 is a multifunctional protein with several well-studied interactions with the host that affect infection ^17^. The transcriptome analysis of IE1 overexpressing cells indicated that the major effects in the context of infection of monocytes are global increase in viral gene expression, which is likely related to the release of repression on the viral genome, and downregulation of interferon response.

IE1 directly binds STAT2, leading to inhibition of ISG expression ^23,40^, which likely explains the downregulation of ISGs observed in cells overexpressing IE1 (Fig. 2b). To test the importance of this interaction, we introduced a deletion that abrogates STAT protein binding ^40,46^ into a Dox-inducible IE1 expression vector and validated that it indeed abolishes the effect of IE1 on ISG expression (Fig. S5a). Nevertheless, overexpression of the STAT-binding mutant IE1 at the time of infection enhanced productive infection to the same extent as the WT IE1 (Fig. 5a), indicating that IE1 does not promote productive infection through downregulation of ISGs or STAT binding. This mutant however partially diminished the effect of IE1 when its expression was induced before infection (Fig. 5b), confirming the role of ISG expression prior to infection in dictating the outcome of infection. Given that IE1-mediated suppression of ISGs does not influence infection outcome, we examined its broader impact. Measurement of infectious progeny showed that cells overexpressing the STAT-binding mutant of IE1 produced only a modest increase relative to control, whereas WT IE1 boosted progeny yield much more efficiently (Fig. 5c). These findings indicate that, although regulation of ISGs by IE1 is dispensable for determining infection outcome, interference with STAT function substantially contributes to infectious progeny production.

**Fig. 5.**
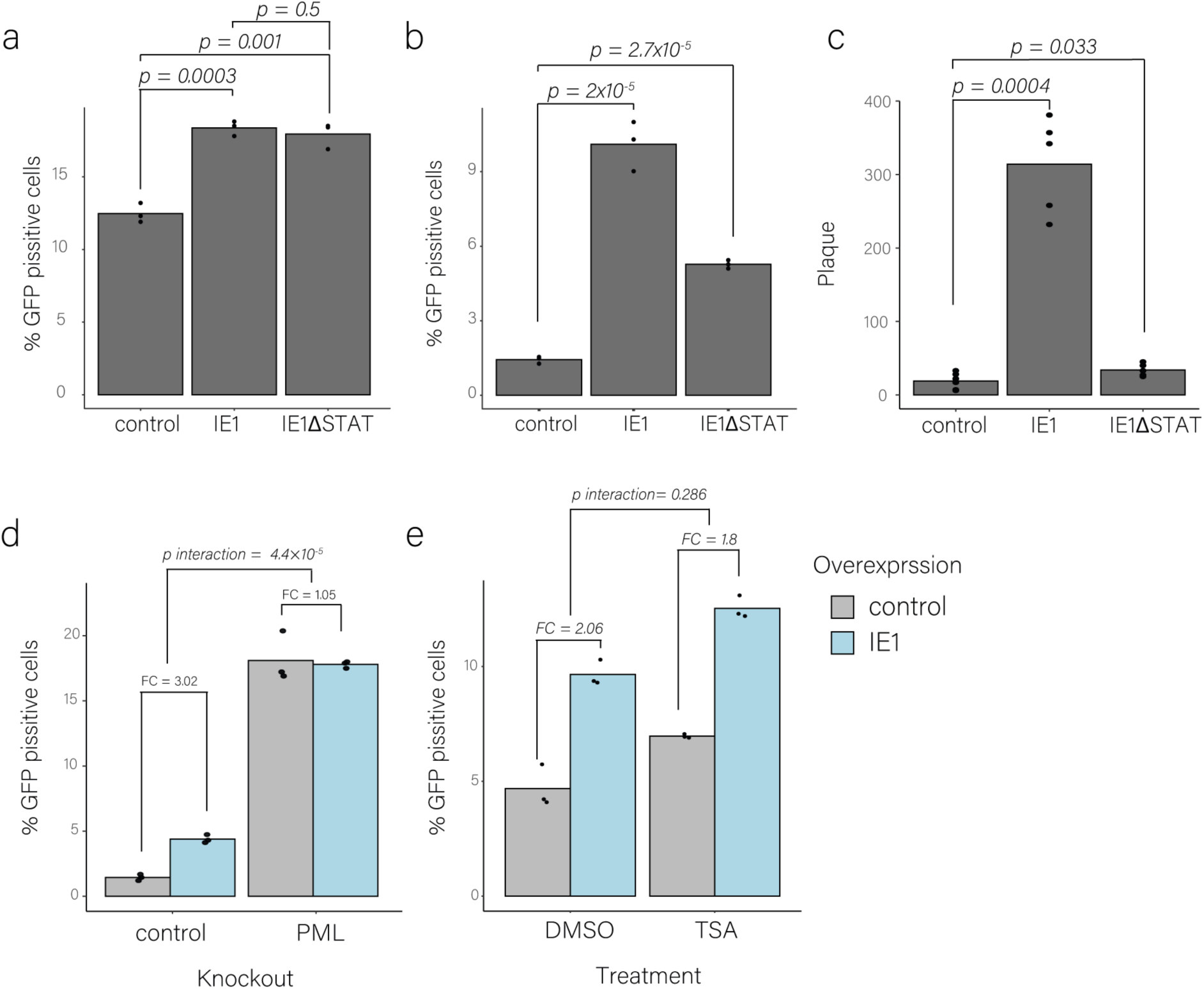
(a). Flow cytometry analysis of THP1-PDGFRa overexpressing control, IE1 and IE1ΔSTAT at the time of HCMV-GFP infection. Cells were analyzed at 3 dpi. P-value was calculated using a two-sided student t-test on logit transformed values. n=3. (b). Flow cytometry analysis of THP1-PDGFRa overexpressing control, IE1 and IE1ΔSTAT 24 hours before HCMV-GFP infection. Cells were analyzed at 3 dpi. p-value was calculated using a two-sided student t-test on logit transformed values. n=3. (c). Measurements of infectious HCMV virus in supernatants collected from infected THP1-PDGFRa overexpressing control, IE1 or IE1ΔSTAT at 10 d.p.i. p-value was calculated using a two-sided student t-test. n=5. (d). Flow cytometry analysis of THP1-PDGFRa with PML or control knockout, overexpressing IE1 and control at the time of HCMV-GFP infection. Cells were analyzed at 3 dpi. P-values were calculated using a two-factor analysis of variance (ANOVA) test on logit transformed values, n=3 (e). Flow cytometry analysis of THP1-PDGFRa treated with TSA or DMSO as a control, overexpressing IE1 and control at the time of HCMV-GFP infection. Cells were analyzed at 3 dpi. P-values were calculated using a two-factor analysis of variance (ANOVA) test on logit transformed values, n=3. P_int_. indicates the P-value of the interaction.

Release of repression on the viral genome, which likely explains the broad enhancement of viral gene expression when IE1 is overexpressed, is related to two known molecular functions of IE1. The first is disruption of PML nuclear bodies ^47^, which are an important determinant of the intrinsic antiviral defense against HCMV infection. The second is direct binding and inhibition of HDAC3 (Nevels et al., 2004), which contributes to deacetylation and silencing of the viral genome. These two functions may also be interconnected since components of the PML bodies are known to bind HDACs ^48^. To test if IE1 effects on infection outcome are mediated by disruption of PML bodies we used CRISPR–Cas9, to generate clonal PML knockout cells (PML KO, Fig. S5b) and tested the effect of IE1 overexpression on infection outcome in these cells. By itself, PML KO significantly promoted productive infection, and strikingly, IE1 expression did not further enhance infection, suggesting that IE1 affected infection outcome mainly by antagonizing PML (Fig. 5d).

We next investigated whether IE1’s impact on infection outcome is also linked to its modulation of histone deacetylation, by examining its effect in cells treated with the HDAC inhibitor (TSA). TSA treatment increased productive infection in THP1 monocytes, but the effect of IE1 was similar to its effect in untreated cells, suggesting that IE1 does not impact infection outcome directly through modulation of histone deacetylation (Fig. 5e). Overall these results illustrate that IE1 main contribution during the establishment of productive infection is to prevent PML association with the viral genome, and the more effective IE1 counteracting PML, the greater the proportion of cells that become productively infected.

**Fig S5.**
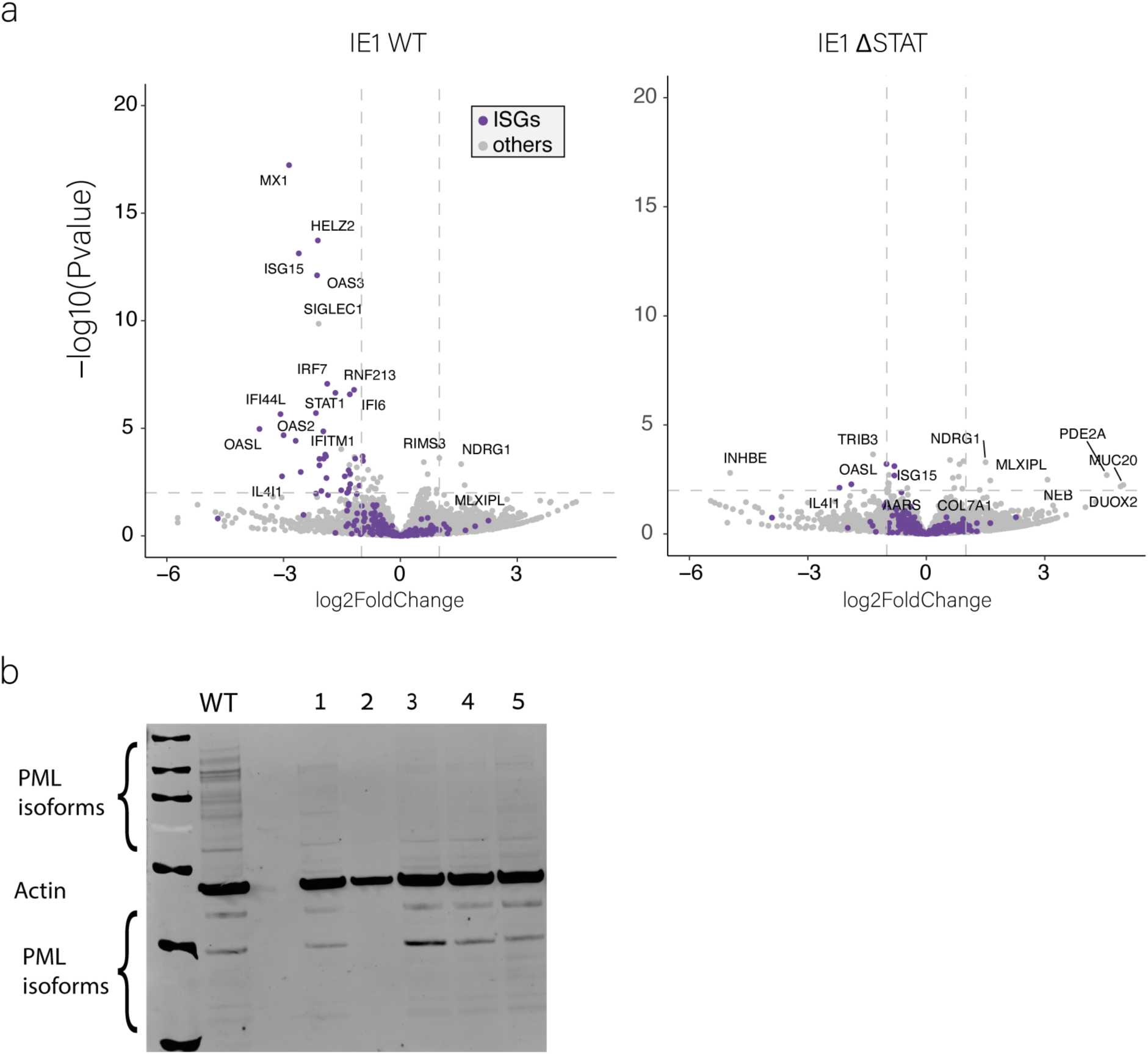
(a) Volcano plot showing the differentially expressed genes of THP1-PDGFRa overexpressing IE1 WT (left) and IE1ΔSTAT (right) compared to the control. Overexpressing cells were induced with Dox for 24 hours prior to RNA extraction and sequencing. Purple dots are interferon-stimulated genes. (b). Western blot analysis of several THP1-PDGFRa clones that were treated with CRISPR-Cas9 to generate Knockout in the PML nuclear bodies. Clone 2 was chosen for the following experiments.

### IE1 promotes reactivation through PML Dissociation

Reactivation from latency is initiated by a trigger that transitions the virus from a repressed state to active gene expression and this process also represents a shift in infection outcome. Expression of IE genes occurs at the onset of reactivation ^49^. To test whether expression of IE genes in latently infected cells serve as a barrier for reactivation, we isolated latently infected THP1 cells by sorting GFP-dim cells, and at 7 dpi, induced the expression of either IE1, IE2 or mCherry as control using Doxycycline and compared it to canonical reactivation by induction of differentiation with PMA. Interestingly, by itself Doxycycline induced reactivation, as measured by emergence of GFP bright cells, likely through induction of stress ^50,51^. Compared to Doxycycline-treated control cells, overexpression of either IE1 or IE2 significantly enhanced reactivation (Fig. 6a), indicating that the levels of expression of either the IE proteins serve as a barrier for reactivation. Simultaneous overexpression of IE1 and IE2 resulted in an additive, but not a synergistic effect, implying IE1 and IE2 contribute to reactivation via distinct, complementary functions.

**Fig. 6.**
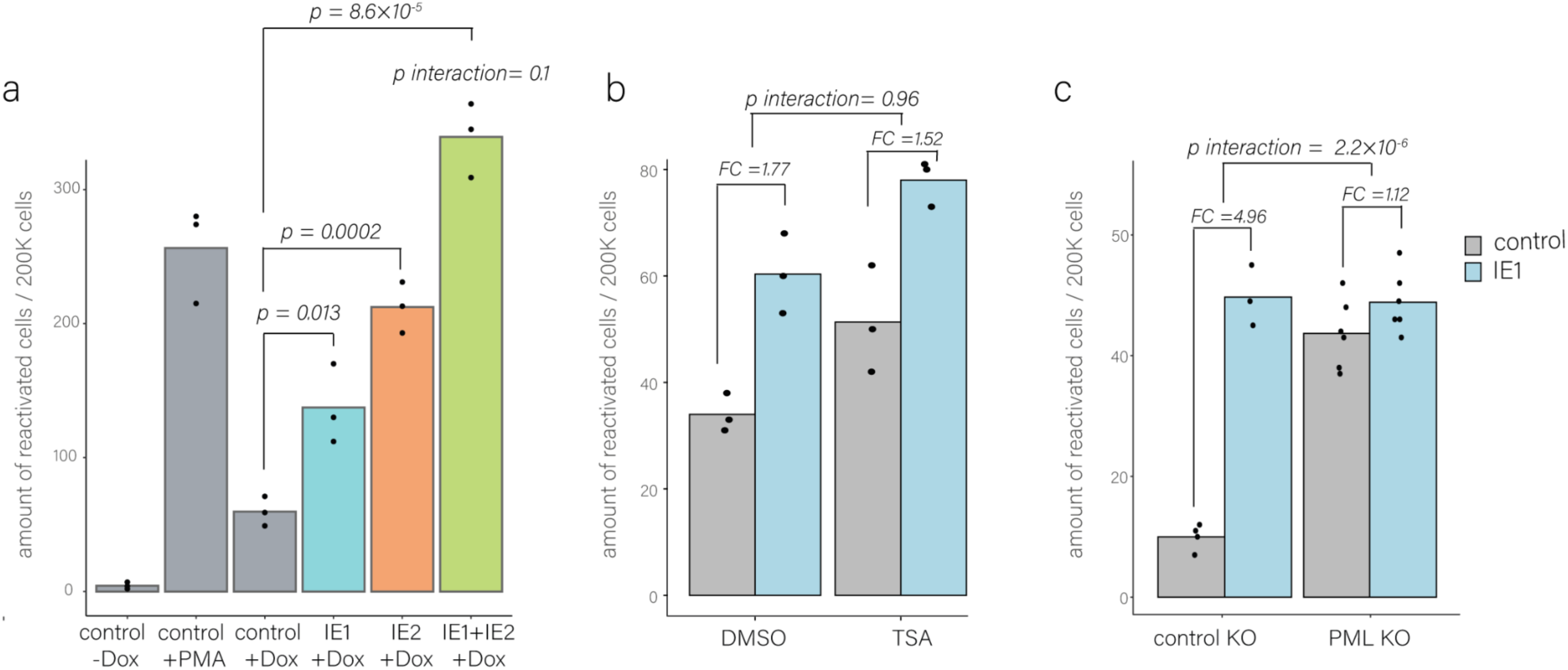
(a). Number of reactivation events in HCMV-infected, GFP-dim sorted, Dox-induced THP1-PDGFRa to overexpress IE1, IE2 and control at 7 d.p.i. Cells were analyzed under the microscope at 3 days post Dox treatment (10 d.p.i.) p-value was calculated using a two-sided student t-test. n=3. (b). Number of reactivation events in HCMV-infected, GFP-dim sorted, Dox-induced THP1-PDGFRa with PML or control Knockout to overexpress IE1 or control at 7 d.p.i. Cells were analyzed under the microscope at 3 days post Dox treatment (10 d.p.i.). P values were calculated using a two-factor analysis of variance (ANOVA) test, n=3 (c). Number of reactivation events in HCMV-infected, GFP-dim sorted, Dox-induced THP1-PDGFRa to overexpress IE1 or control at 7 d.p.i alongside treatment with TSA or DMSO control at the time of induction. Cells were analyzed under the microscope at 3 days post Dox treatment (10 d.p.i.). P values were calculated using a two-factor analysis of variance (ANOVA), n = 3-6. P_int_. indicates the P-value of the interaction.

We further dissected which of IE1’s functions is involved in reactivation. Reactivation following induction of either WT IE1 or STAT-binding mutant IE1 was comparable, indicating that the modulation of ISG expression by IE1 does not affect its ability to induce reactivation (Fig. S6).

We next assessed the contribution of IE1’s modulation of histone deacetylation. As previously reported ^29,52^, TSA treatment induced reactivation in infected THP1 monocytes, however the effect of IE1 on reactivation was comparable between TSA treated and untreated cells, suggesting that IE1 likely promotes reactivation independent of its modulation of histone deacetylation (Fig. 6b). To determine whether the ability of IE1 to promote reactivation is linked to its function in disrupting PML nuclear bodies, we measured reactivation in either control or PML KO THP1 monocytes, infected with HCMV and sorted for non-productive cells. PML deficiency and IE1 overexpression promoted reactivation to a similar extent, and IE1 overexpression did not further increase reactivation in PML-deficient cells (Fig. 6c). These results indicate that the enhancement of reactivation by IE1 is primarily through counteracting the activity of PML, suggesting that PML has an active role in the maintenance of at least early stages of latency.

Overall, these results illustrate the key role of HCMV IE proteins in determining the outcome of infection. Moreover, it shows that expression of either IE1 or IE2 and to a greater extent the expression of both, promotes reactivation in latent monocytes.

**Fig. S6.**
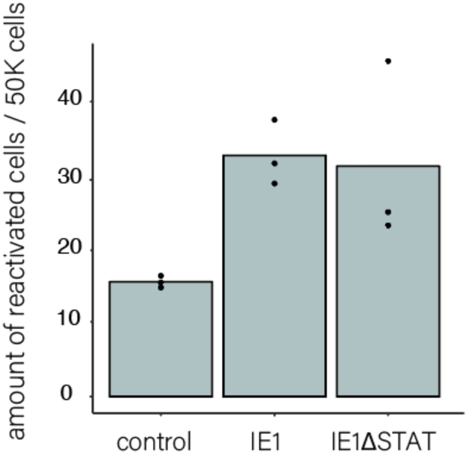
Number of reactivation events in HCMV-infected, GFP-dim sorted, Dox-induced THP1-PDGFRa to overexpress IE1, IE1ΔSTAT and control at 7 d.p.i. Cells were analyzed under the microscope at 3 days post Dox treatment (10 d.p.i.).

## Discussion

Expression of HCMV immediate early (IE) genes is a central determinant in the balance between lytic and latent infection. Chromatin-mediated repression of IE gene expression is a hallmark of latent infection, observed both in experimental systems and in natural infection ^25–28^. Although regulation of IE gene expression is considered a key aspect of HCMV latency and reactivation, consistent with previous findings ^34^, we show here that ectopic expression of IE genes alone is insufficient to trigger productive infection in THP1 cells, indicating an additional infection block in monocytes ^34^. We recently demonstrated that this block largely reflects inefficient entry ^36^. Here, we show that when entry is enhanced by overexpression of the HCMV entry receptor PDGFRα, ectopic IE gene expression markedly increases productive infection. This system therefore provides a framework to dissect the specific functions of IE genes that promote lytic infection.

Transcriptome analysis of IE1 and IE2 in infected and uninfected cells provided insights into their mechanisms of action in shaping infection outcome in monocytes. IE1 overexpression led to a global increase in viral gene expression and its ability to promote lytic infection predominantly was found to depend primarily on disruption of PML nuclear bodies. These findings indicate that IE1 facilitates HCMV lytic infection by relieving PML-mediated repression of viral transcription. In contrast, while histone deacetylase inhibition on its own influences infection outcome, this activity of IE1 makes little contribution to its capacity to promote lytic infection. Another prominent effect of IE1 overexpression was suppression of the interferon response pathway, however this function did not influence infection outcome, consistent with our previous observations that ISG induction after infection, does not affect infection outcome ^33^. Instead, ISG levels at the time of infection are critical for determining the infection outcome, which likely explains why IE1 expression prior to infection exerts a stronger effect than induction at or after infection. Although STAT binding-mediated inhibition of ISG expression by IE1, does not determine infection outcome, it contributes substantially to later stages of infection, enhancing the production of infectious progeny. Notably, the ability of IE1 to promote reactivation is also mainly dependent on its PML disruption function, suggesting it further has a role in latency maintenance ^53^.

Unlike IE1, IE2 does not increase global viral gene expression at early stages of infection, rather selectively upregulates early genes while downregulating a subset of true-late genes, consistent with its established direct effect on viral gene expression ^54,55^.

IE2 also affected ISG expression, but in a context-dependent manner. In uninfected cells, IE2 robustly induced interferon-stimulated genes (ISGs), possibly as a consequence of DNA damage ^43^. In contrast, induction of IE2 at the time of infection did not elicit the same antiviral response, suggesting that additional viral factors, perhaps IE1, mitigate its otherwise toxic effects. This aligns with previous observations that cytotoxicity of IE2 overexpression in uninfected cells can be alleviated when its expression is induced only in infected cells ^56^. IE2 also exerted distinct effects on the cell cycle in infected and uninfected cells. In infected cells, the upregulation of E2F targets and G2/M checkpoint pathways are consistent with IE2 pushing the cells towards S phase ^57^. Since these changes were absent when IE2 is induced prior to infection, and in these settings IE2 did not promote lytic infection, this suggests that IE2’s effect on the cell cycle may contribute to its role in determining infection outcome.

Intriguingly, induction of IE2 expression, enhanced reactivation in latent cells, suggesting it is capable of overcoming the repressive structures existing in latent cells to induce the expression of viral genes. Further studies are needed to understand the molecular mechanisms underlying this transcriptional induction.

In summary, we show that both IE1 and IE2 expression serve as critical barriers to productive HCMV infection. For IE1, this effect is mediated through disruption of PML bodies, which likely relieves repression of the viral genome and thereby promotes broad viral gene expression. For IE2, the phenotype is likely linked to its capacity to drive early gene expression, although additional work is required to establish a direct causal mechanism.

## Material and Methods

### Cell culture and virus

293T cells (ATCC CRL-3216) and Primary human foreskin fibroblasts (ATCC CRL-1634) were maintained in DMEM with 10% FBS, 2 mM L-glutamine and 100 units ml−1 penicillin and streptomycin (Beit-Haemek).

THP1 cells, purchased from ATCC (TIB-202) were grown in RPMI media with 20% heat-inactivated FBS, 2 mM L-glutamine and 100 units/ml penicillin and streptomycin (Beit-Haemek).

TB40E strain of HCMV, containing an SV40–GFP reporter ^35^ was used for all the experiments. Virus propagation was done by adenofection of a bacterial artificial chromosome of the viral genome into fibroblasts as described previously (Elbasani et al., 2014). When most of the cells in the culture died, supernatant was collected and cleared from cell debris by centrifugation.

### Infection procedures

24h Before infection, cells were grown in 0.5% for 24h. Infection was performed by centrifugation at 800g for 1h in 24-well with the virus added at multiplicity of infection (MOI) = 5, followed by washing and supplementing with fresh media. Notably, because this MOI is based on quantification of infectious particles in fibroblasts it is effectively lower in monocytic cells.

Cells were treated with 1 µg/ml Doxycycline (Dox) at the indicated time points before or during infection. For experiments analyzed at 3 dpi, cells were washed at 1 hpi to remove residual virus and then supplemented with fresh medium containing Dox. Dox was completely removed at 24 hpi by washing the cells. Where indicated, 1 µM TSA or DMSO (vehicle control) was added either at the time of infection or at the time of reactivation stimulation.

For progeny assay, at 10 dpi the supernatant was cleared from cell debris by centrifugation and transferred to fibroblasts. Infected fibroblasts were counted 2-3 days later.

For Reactivation assays, GFP-dim cells were sorted at 5 dpi and at 7 dpi were treated with the indicated stimulators. HCMV-positive cells were counted on a fluorescent microscope at 10 dpi.

### Flow cytometry and sorting

Cells were analyzed on a BD Accuri C6 or CytoFLEX (Beckman Coulter) and sorted on a BD FACS AriaIII using FACSDiva software. All analyses and figures were done with FlowJo.

### Next generation sequencing

RNA-seq library preparation was performed as described previously ^58^. Cells were collected with Tri-Reagent (Sigma-Aldrich). Total RNA was extracted according to the manufacturer instructions and poly-A selection was performed using Dynabeads mRNA DIRECT Purification Kit (Invitrogen). The mRNA samples were subjected to DNase I treatment and 3ʹ dephosphorylation using FastAP Thermosensitive Alkaline Phosphatase (Thermo Scientific) and T4 polynucleotide kinase (New England Biolabs) followed by 3ʹ adaptor ligation using T4 ligase (New England Biolabs). The ligated products were used for reverse transcription with SSIII (Invitrogen) for first-strand cDNA synthesis. The cDNA products were 3ʹ-ligated with a second adaptor using T4 ligase and amplified for 8 cycles in a PCR for final library products of 200–300 bp. Raw sequences were first trimmed at their 3’ end, removing the Illumina adapter and poly(A) tail. Alignment was performed using Bowtie 1 ^59^ (allowing up to two mismatches) and reads were aligned to the human (hg19) or TB40E (NCBI EF999921.1) strain of HCMV. Reads aligned to ribosomal RNA were removed. Reads that were not aligned to the genome were then aligned to the transcriptome.

### Differential expression and enrichment analysis

Differential expression analysis on RNA-seq data was performed with DESeq2 (v.1.22.2) using default parameters, with the number of reads in each of the samples as an input.

The log2(fold change) values from the DE on the RNA-seq was used for enrichment analysis using GSEA (v.4.1). For the analysis, only genes with a minimum of ten reads were included.

Gene list of cellular ISGs compiled based on ^60^ and gene list of cell cycle -related genes was based on compiled list of the pathways E2F targets, MYC targets and G2M checkpoints from hallmark.

### Immunofluorescence

For HCMV replication compartment detection, cells were plated on i-bidi slides, fixed in 4% paraformaldehyde for 15 min, washed in PBS, permeabilized with 0.1% Triton X-100 in PBS for 10 min and then blocked with 10% goat serum in PBS for 30 min. Immunostaining was performed for the detection of mouse anti-UL44 (CA006-100, Virusys) with 2% goat serum diluted in PBS. Cells were washed 3 times with PBS and labeled with goat anti-mouse–Alexa Fluor 647 (Thermo Fisher) secondary antibody and DAPI (4ʹ,6-diamidino-2-phenylindole) diluted 1:500 in PBS for 1 hour at room temperature, followed by 3 PBS washes.

### Plasmid construction and lentiviral transduction

PDGFRα plasmid was cloned as described (my paper). sgRNA plasmids to knockout PML were cloned into with lentiCRISPR v2 plasmid (Addgene no. 52961 ^61^). To generate inducible expression plasmids of IE1 and IE2, genes were amplified from cDNA of infected cells and validated by Sanger sequencing. IE1 with STAT mutation (IE1ΔSTAT) as described in (IE1dl410-420,^46^) was ordered from TWIST. The genes were cloned into pLVX-Puro-TetONE-SARS-CoV-2-nsp1-2XStrep (kind gift from N. Krogan, UCSF) in place of the SARS-CoV-2-nsp1-2XStrep cassette using linearization with BamHI and EcoRI (NEB). The genes were amplified with primers containing flanking regions homologous to the vector (Supplementary Table 1). The amplified PCR fragments were cleaned using a gel extraction kit (QIAGEN) according to the manufacturer’s protocol and were cloned into the vectors using a Gibson assembly reaction (NEB). Inducible mCherry, cloned on the same backbone was previously described^33^ and was used as control in all experiments with these inducible expression plasmids.

Lentiviral particles were generated by cotransfection of the expression constructs and second-generation packaging plasmids (psPAX2, Addgene, catalog no. 12260 and pMD2.G, Addgene, catalog no. 12259), using jetPEI DNA transfection reagent (Polyplus transfection) into 293T cells, according to the manufacturer’s instructions. At 60 h post-transfection, supernatants were collected and filtered through a 0.45-μm polyvinylidene difluoride filter (Millex). THP1 cells were transduced with lentiviral particles by centrifugation at 800g for 1h in 24-well plates. 2 days post transfection the cells were transferred to selection media (blasticidin, 10 μg/ ml for 5 days or puromycin, 1.75 μg/ml for 4 days). Blasticidin and puromycin were removed and cells were recovered for at least two days before subsequent processing.

### Quantitative real-time PCR analysis

Total DNA was extracted using QIAamp DNA Blood kit (Qiagen) according to the manufacturer’s instructions. Real-time PCR was performed using SYBR Green PCR master-mix (ABI) on the QuantStudio 12K Flex (ABI). Amplification of NRP2 and ITGB3 was normalized to the host gene ANXA5 (primers detailed in table 1). Amplification of the viral gene UL44 was normalized to the host gene B2M (primer detailed in table 1).

**Table 1.**
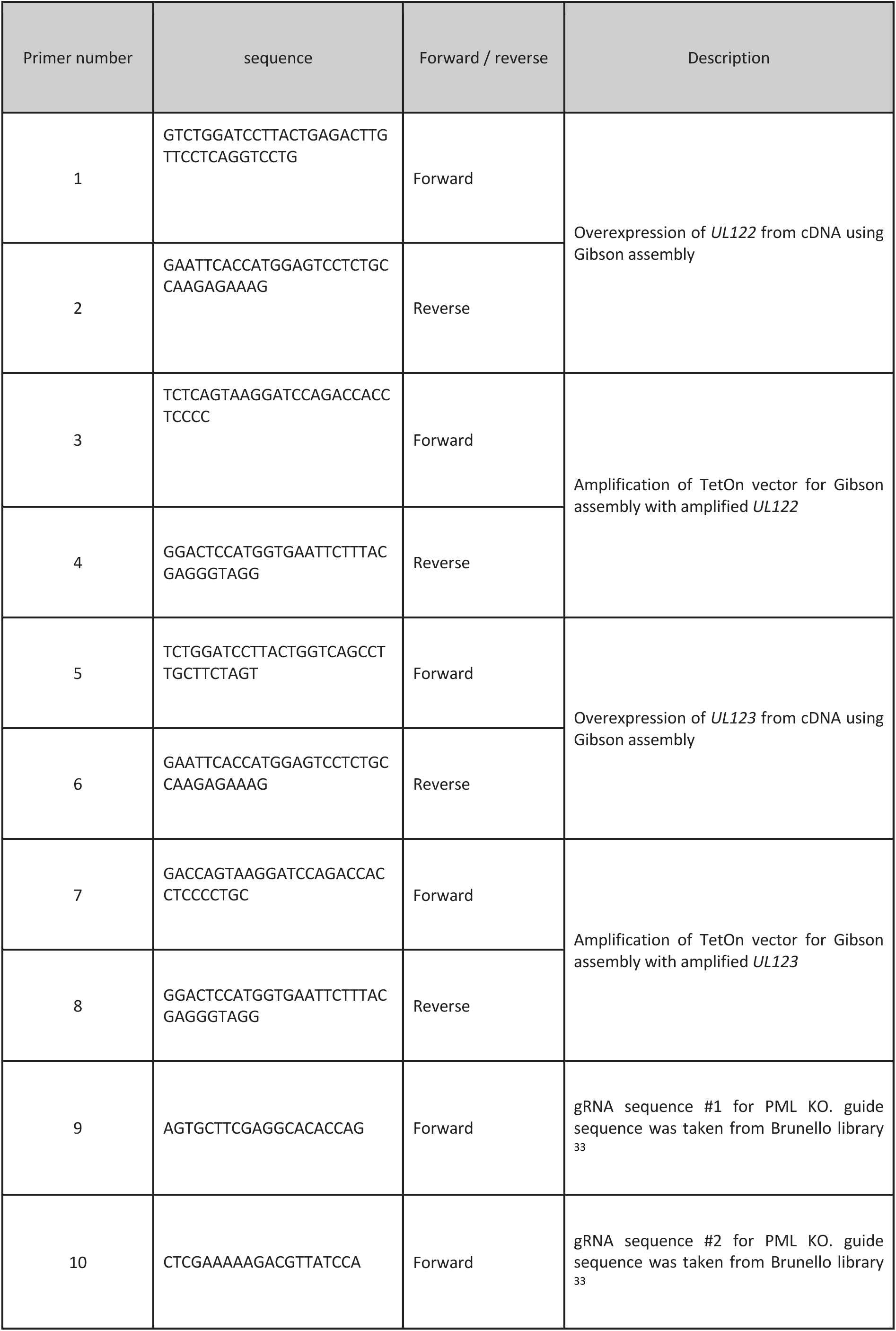
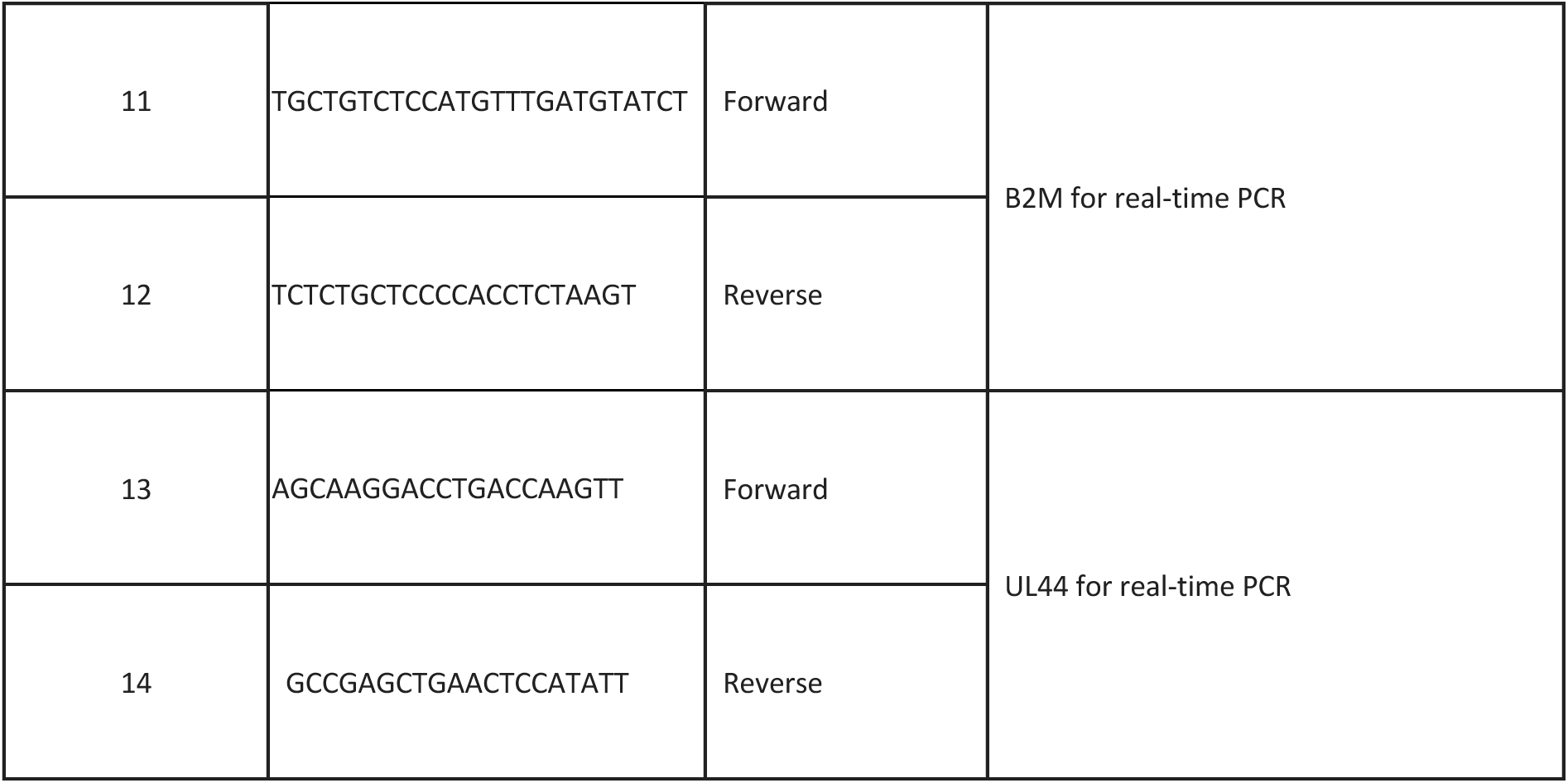
primers list.

### Western blot

For western blot analysis, cells were lysed in RIPA buffer (150 mM NaCl,1% NP-40, 0.5% sodium deoxycholate, 0.1%SDS, 50 mM Tris-HCl pH 7.4, and 1×EDTA-free protease inhibitor cocktail). Lysates were cleared by centrifugation and supplemented with Sample Buffer. Proteins were separated by SDS-PAGE electrophoresis, transferred to nitro-cellulose membranes (0.25 mm, ThermoFisherScientific), and detected using an infrared fluorescent antibody detection system (LI-COR) using the antibodies IE (abcam, CH160), mCherry (abcam, ab167453), Actin (Sigma, A4700), PML (Santa Cruz, sc-377340) and ISG15 (cell signaling, 2743s) diluted 1:1000. The secondary antibodies were goat anti-rabbit–IRDye 800CW and goat anti-mouse–IRDye 680RD (LI-COR Biosciences) diluted 1:7,500. Membranes were visualized in a LI-COR Biosciences Odyssey imaging system.

## Data availability

All next-generation sequencing data files have been deposited in Gene Expression Omnibus under accession number GSE307171.

## Acknowledgments

We thank the members of the Stern-Ginossar lab for the critical reading of the manuscript. We would like to thank the Weizmann flow cytometry and microscopy unit for technical assistance. This study was supported by a European Research Council consolidator grant (CoG-2019-864012) and an Israel Science Foundation grant to N.S.-G. (2507/23).

## Author contributions

Y.K., N.S.-G. and M.S. conceived and designed the project. Y. K., and M.S. performed the experiments. Y.K., A.N., N.S-G and M.S. analyzed and interpreted the data. Y.K., N.S.-G. and M.S. wrote the manuscript with input from all the authors.

